# Type I IFN expression is inhibited during cell division by CDK4/6

**DOI:** 10.1101/2023.04.15.537049

**Authors:** Rebecca P. Sumner, Ailish Ellis, Sian Lant, Hannah Ashby, Greg J. Towers, Carlos Maluquer de Motes

## Abstract

Cells are equipped to defend themselves from invading pathogens through sensors such as cGAS, which upon binding DNA induces type I interferon (IFN) expression. Whilst IFNs are crucial for limiting viral infection and activating adaptive immunity, uncontrolled production causes excessive inflammation and autoimmunity. cGAS binds DNA of both pathogenic and cellular origin and its activity is therefore tightly regulated. This is particularly apparent during mitosis, where cGAS association with chromatin following nuclear membrane dissolution and phosphorylation by mitotic kinases negatively regulate enzymatic activity. Here we describe a novel mechanism by which DNA sensing and other innate immune pathways are regulated during cell division, dependent on cyclin dependent kinases (CDK) 4 and 6. Inhibition of CDK4/6 using chemical inhibitors, shRNA-mediated depletion, or overexpression of cellular CDK4/6 inhibitor p16INK4a, greatly enhanced DNA- or cGAMP-induced expression of cytokines and IFN-stimulated genes (ISG). Mechanistically, CDK4/6-dependent inhibition mapped downstream of cytoplasmic signalling events including STING and IRF3 phosphorylation, limiting innate immune induction at the level of IFNβ mRNA expression. This regulation was universal, occurring in primary and transformed cells of human and murine origin, and broad, as IFNβ expression was inhibited in a CDK4/6-dependent manner downstream of multiple pattern recognition receptors. Together these findings demonstrate that host innate responses are limited by multiple mechanisms during cell division, thus defining cellular replication as an innate immune privileged process that may be necessary to avoid aberrant self-recognition and autoimmunity.

## Introduction

Innate immunity constitutes an important first line of defence against invading pathogens carrying pathogen-associated molecular patterns (PAMPs), as well as against inducers of damage-associated molecular patterns (DAMPs) such as tumorigenesis. Engagement of PAMPs and DAMPs with pattern recognition receptors (PRRs) induces complex signalling cascades, culminating in the activation of transcription factors of the interferon (IFN) regulatory factor (IRF) and nuclear factor kappa B (NF-κB) families and subsequent expression of IFNs and pro-inflammatory cytokines and chemokines(Li & Wu, 2021). Binding of IFNs to IFN receptors activates signalling cascades dependent on the Janus kinase (JAK) and signal transducer and activator of transcription (STAT) proteins, inducing the expression of hundreds of IFN-stimulated genes (ISGs) (Hu *et al*, 2021). Whilst some of these proteins directly block pathogen replication, or induce apoptosis, the inflammatory milieu is also crucial for attracting antigen presenting cells and activating cells of the adaptive arm of the immune system. PRRs include toll like receptors (TLRs) that sense nucleic acids and microbial membrane components such as lipids and proteins, the RIG-I-like receptors (RLRs) that sense various forms of pathogen RNA, DNA sensors such as the AIM2-like receptors (ALRs) and cGAS, and the NOD-like receptors (NLRs), which sense a variety of PAMPs and DAMPs including nucleic acids, proteins and metabolites and result in inflammasome activation(Li & Wu, 2021; Zahid *et al*, 2020). Uncontrolled or aberrant activation of these receptors leads to excessive IFN and inflammatory cytokine production and underlies many autoimmune conditions(Okude *et al*, 2020).

The major sensor for pathogenic DNA in the cell is cyclic GMP-AMP synthase (cGAS), an enzyme that catalyses the formation of cyclic dinucleotide 2’3’ cyclic GMP-AMP (cGAMP) from ATP and GTP(Ablasser *et al*, 2013; Sun *et al*, 2013). cGAMP binds and activates ER-resident stimulator of IFN genes (STING), which involves its dimerisation, phosphorylation and translocation to the Golgi where it recruits tank-binding kinase 1 (TBK1) and IRF3(Tanaka & Chen, 2012; Wu *et al*, 2013). TBK1 then phosphorylates IRF3, inducing nuclear translocation and the expression of IRF-dependent genes including type I IFN and ISGs. cGAMP binding to STING also induces the activation of NF-κB, dependent on kinases TAK1 and the IKK complex(Balka *et al*, 2020; Yum *et al*, 2021). The binding of dsDNA to cGAS induces its oligomerisation, a prerequisite for enzymatic activity(Li *et al*, 2013; Zhang *et al*, 2014), and therefore cGAS sensing is dependent on DNA length(Luecke *et al*, 2017). DNA binding is however sequence independent, with interactions observed between cGAS and the phosphate-sugar backbone along the minor groove(Civril *et al*, 2013), raising the question of how cGAS distinguishes between self and non-self. Indeed, cGAS is competent to bind DNA of cellular origin, including mitochondrial DNA(Huang *et al*, 2020), micronuclei(Mackenzie *et al*, 2017), damaged/leaked nuclear DNA(Zhou *et al*, 2021) and engulfed cellular debris from dying cells(King *et al*, 2017). To avoid aberrant recognition, cGAS activity is subject to complex regulation by phosphorylation, SUMOylation, glutamylation, ubiquitination and protein-protein interactions(Motwani *et al*, 2019).

Cell division, ending in mitosis, is regulated by a coordinated cascade of cyclin expression that stimulates the activity of the cyclin-dependent kinases (CDKs), which in turn regulate the transition from one cell cycle stage to the next(Malumbres, 2014). How DNA sensing is regulated during this fundamental process, which involves nuclear membrane breakdown, has remained somewhat elusive. Recent publications from the Chen and Shu labs have however now begun to shed some light on this. Firstly, cGAS association with chromatin following nuclear envelope breakdown prevents its oligomerisation and activation(Li *et al*, 2021). Secondly, cGAS is inactivated through phosphorylation by mitotic kinases such as Aurora kinase B(Li *et al.*, 2021) and CDK1(Zhong *et al*, 2020), both of which block cGAMP production. We hypothesised that multiple mechanisms would likely exist to regulate DNA sensing during cell division, so we blocked cell cycle at various stages using CDK inhibitors and assessed the ability of cells to respond to stimulation. We found that inhibition of CDK4/6 activity, either chemically or genetically, led to a significantly enhanced DNA sensing response and ISG expression in primary and transformed cells of both human and murine origin. This regulation mapped downstream of STING and IRF3 activation, at the level of type I IFN expression. Consistent with downstream regulation, CDK4/6 dampened innate activation induced by multiple PRRs, including TLR4 and the RLRs. CDK4/6-mediated regulation of innate immunity was also independent of the previously described inflammatory induction associated with cell cycle arrest, senescence and endogenous retrovirus (ERV) reactivation following long-term CDK4/6 inhibition(Gluck *et al*, 2017; Goel *et al*, 2017). Together this study identifies a novel and direct role for CDK4/6 in negatively regulating type I IFN expression, further defining cell division as an immune privileged process that may be critical to avoid aberrant self-sensing and autoimmune induction.

## Results

### CDK4/6 inhibitors enhance DNA sensing-dependent ISG induction

Hypothesising that multiple mechanisms would exist to regulate DNA sensing during cell division to prevent recognition of cellular patterns, we inhibited the cell cycle with CDK inhibitors (Fig 1A) and measured subsequent DNA sensing responses. Innate induction was assessed by measuring luciferase activity in the supernatants of monocytic THP-1 cells that had been modified to express Gaussia luciferase under the control of the IFIT-1 (also known as ISG56) promoter, which is both IRF3- and IFN-sensitive(Mankan *et al*, 2014). All three inhibitors arrested cellular replication at the doses tested, as expected (data not shown). Whilst inhibition of CDK1 with RO-3306 or CDK2 with K03861 had minimal effect on IFIT-1 reporter activity following stimulation of cells with DNA sensing agonists herring testis DNA (HT-DNA) or cGAMP, inhibition of CDK4/6 with palbociclib greatly enhanced innate activation in response to both agonists (Fig 1B). This finding was confirmed by measuring cGAMP-induced endogenous expression of classical ISGs, such as *CXCL-10*, *IFIT-2* and *MxA*, by qPCR (Fig 1C) and CXCL-10 protein expression by ELISA (Fig 1D). A similar effect on DNA sensing-induced reporter activity was also observed with another CDK4/6 inhibitor, ribociclib (Fig 1E). As expected, both palbociclib and ribociclib increased the proportion of cells in the G0/G1 phase of the cell cycle, as CDK4/6 activity is required for progression from G1 to S phase (Suppl. Fig 1A).

**Fig 1:**
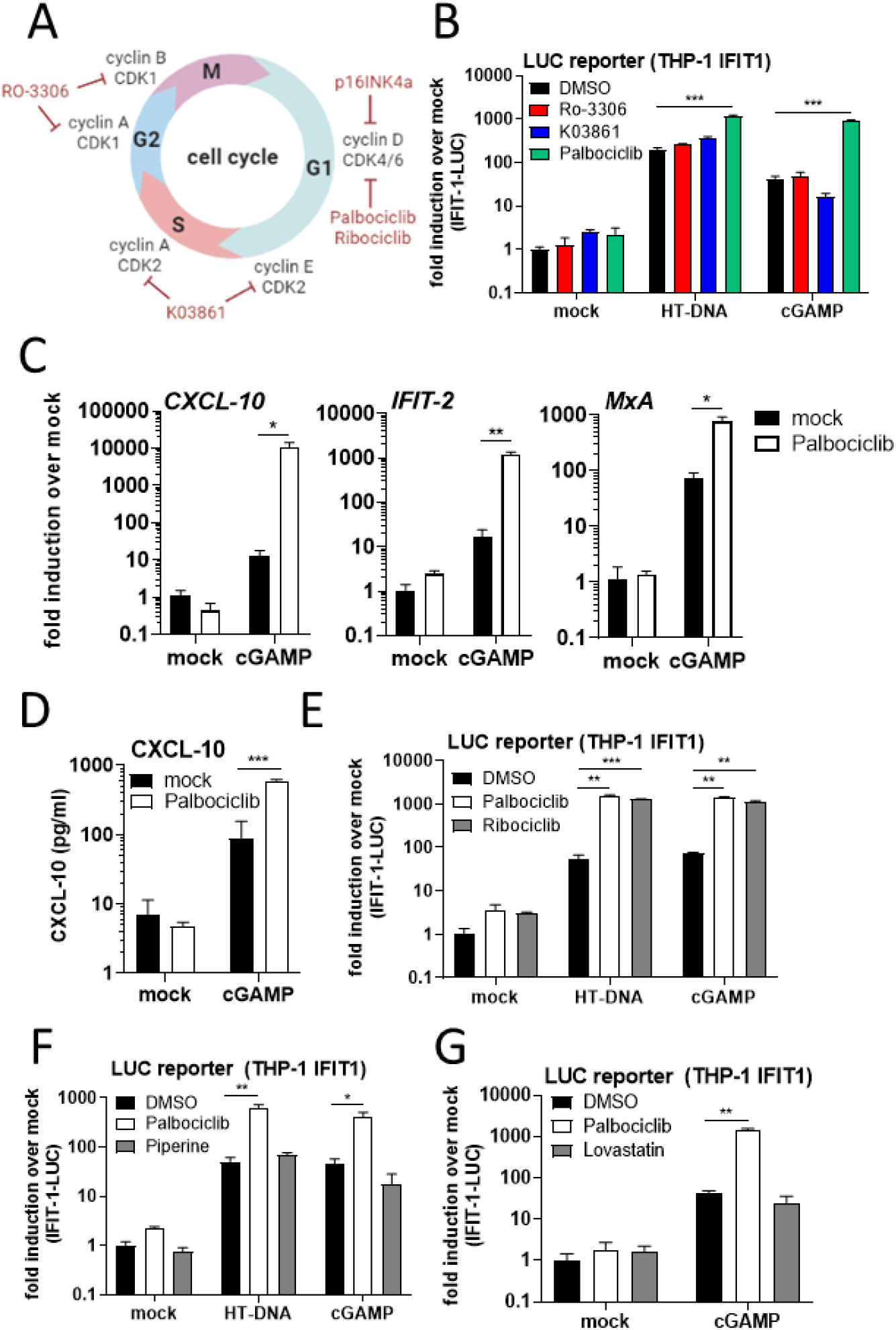
CDK4/6 inhibition enhances DNA sensing-dependent ISG induction. (A) Cell cycle graphic. Created with BioRender.com. (B) Luciferase activity from THP-1 IFIT-1 cells arrested for 48 h with inhibitors of CDK1 (RO-3306, 1 μM), CDK2 (K03861, 0.5 μM), CDK4/6 (palbociclib, 2 μM) or DMSO control and stimulated for 16 h by transfection with HT-DNA (50 ng/ml) or cGAMP (500 ng/ml), or mock treated. (C) ISG qRT-PCR from THP-1 IFIT-1 cells mock-treated or treated for 48 h with 2 μM palbociclib and stimulated overnight by transfection with cGAMP (500 ng/ml). (D) CXCL-10 in the supernatants from (C), measured by ELISA. (E-G) Luciferase activity from THP-1 IFIT-1 cells arrested for 48 h with palbociclib (2 μM, E-G), ribociclib (2 μM, E), piperine (25 μM, F), lovastatin (10 μM, G) or DMSO control and stimulated overnight by transfection with HT-DNA (50 ng/ml) or cGAMP (500 ng/ml), or a mock-treated control. Data are mean ± SD, n = 3, representative of at least 3 repeats. Fold inductions were calculated by normalising each condition with the non-drug treated, non-stimulated (mock) control. Statistical analyses were performed using Student’s t-test, with Welch’s correction where appropriate. *P < 0.05, **P < 0.01, ***P < 0.001.

### CDK4/6 inhibitor-mediated regulation of DNA sensing is independent of cell cycle arrest and senescence

To determine whether enhanced ISG expression was specific to CDK4/6 inhibition or was a general phenomenon of G1 arrest, we treated cells with other small molecules that had been documented to induce G1 arrest in a CDK4/6-independent manner(Fofaria *et al*, 2014; Rao *et al*, 1999). Whilst both piperine (Suppl. Fig 1B) and lovastatin (Suppl. Fig. 1C) increased the proportion of cells in G0/G1, neither enhanced ISG induction in response to DNA sensing agonists (Fig 1F, G) suggesting innate immune regulation was specific to CDK4/6 inhibition. Long-term treatment with CDK4/6 inhibitors induces cell cycle arrest and replication stress(Crozier *et al*, 2022), as well as senescence and inflammatory cytokine expression(Gluck *et al.*, 2017; Yoshida *et al*, 2016), however enhanced ISG expression was observed after short palbociclib pre-treatment (7h, Suppl. Fig 1D), when G1 arrest (Suppl. Fig 1E), nor markers of senescence such as loss of MCM2(Harada *et al*, 2008), were observed (Suppl. Fig 1F). As expected, reduced expression of MCM2 was observed following 48 h treatment of THP-1 cells with DNA damage inducer doxorubicin (Suppl. Fig 1F). Together these results indicated that CDK4/6 inhibitor-mediated regulation of DNA sensing responses was independent of cell cycle arrest and senescence.

### CDK4/6 negatively regulate DNA sensing

To determine a specific role for CDK4/6 in palbociclib/ribociclib-mediated enhancement of DNA sensing responses we undertook a genetic approach to manipulate CDK4/6 activity. Firstly, we generated a lentiviral vector overexpressing a FLAG-tagged version of cellular CDK4/6 inhibitor p16INK4a(McConnell *et al*, 1999), or FLAG-GFP as a negative control. THP-1 cells transduced with these vectors expressed detectable levels of FLAG-tagged GFP and p16INK4a (Fig 2A), and G1 arrest was observed with the p16INK4a-, but not GFP-expressing lentivector, as expected (Suppl. Fig 2). When transduced cells were stimulated by transfection with HT-DNA or cGAMP, IFIT-1 reporter activity was significantly enhanced in the p16INK4a-expressing cells compared to GFP-expressing controls, confirming data obtained with CDK4/6 inhibitors. Furthermore, we delivered lentiviral vectors expressing CDK-specific shRNAs to THP-1 cells and found that depletion of CDK6 also augmented DNA sensing-induced IFIT-1 reporter activity (Fig 2C). Interestingly targeting CDK4 did not boost ISG expression in these experiments, however this may have been due to incomplete depletion of CDK4 expression (Fig 2D) that did not result in an increased proportion of cells in G0/G1, as was observed with CDK6 depletion (Suppl. Fig 2B).

**Fig 2:**
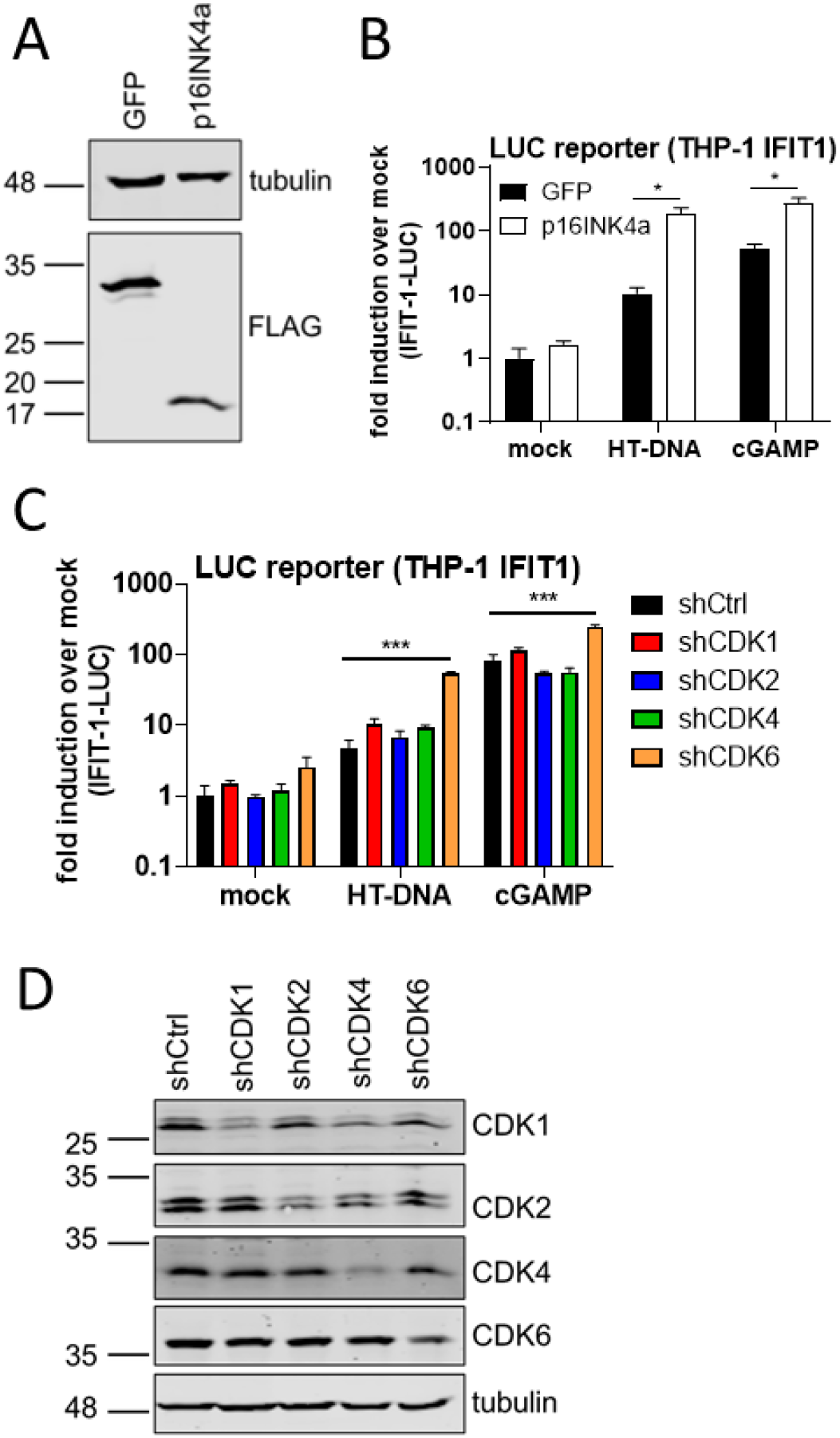
CDK4/6 negatively regulate DNA sensing. (A) Immunoblot of THP-1 IFIT-1 cells transduced for 48 h with FLAG-GFP- or FLAG-p16INK4a-expressing lentiviruses probed for FLAG and tubulin. (B) Luciferase activity from cells from (A), stimulated overnight by HT-DNA (50 ng/ml) or cGAMP (500 ng/ml) transfection. (C) Luciferase activity from THP-1 IFIT-1 cells transduced for 48 h with lentiviruses expressing shRNA against CDKs or an shCtrl and stimulated overnight by transfection with HT-DNA (50 ng/ml) or cGAMP (500 ng/ml). (D) Immunoblot from (C), probed for CDK1, CDK2, CDK4, CDK6 and tubulin. Data are mean ± SD, n = 3, representative of at least 3 repeats. Fold inductions were calculated by normalising each condition with the FLAG-GFP transduced, non-stimulated (mock) control (B) or shCtrl transduced, non-stimulated (mock) control (D). Statistical analyses were performed using Student’s t-test, with Welch’s correction where appropriate. *P < 0.05, ***P < 0.001.

To investigate whether CDK4/6 regulated DNA sensing in primary cells we treated human foetal foreskin fibroblasts (HFFF) with palbociclib (Suppl. Fig 3A), ribociclib (Suppl. Fig 3B) or transduced them with the p16INK4a-expressing lentivector (Suppl. Fig 3C) and in all cases observed a significant increase in DNA sensing-dependent ISG expression. Enhanced ISG induction was also apparent in A549 (Suppl. Fig 3D), U2OS (Suppl. Fig 3E) and murine fibroblasts (McCoy cells, Suppl. Fig. 3F) treated with palbociclib and stimulated with HT-DNA/cGAMP. Taken together, these data define a universal role for CDK4/6 in negatively regulating DNA sensing-induced ISG expression in both primary and transformed cells of human and murine origin.

**Fig 3:**
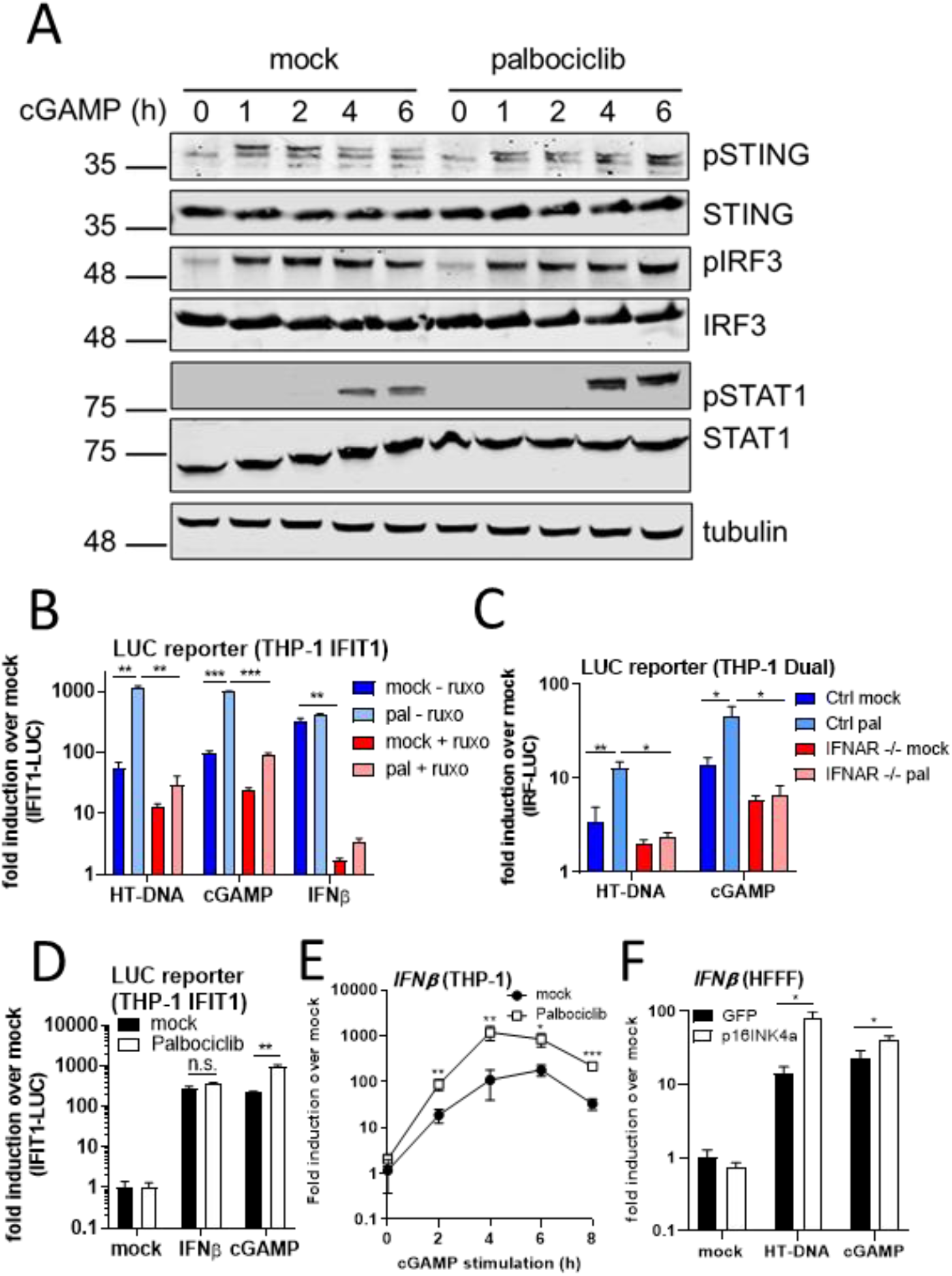
CDK4/6 negatively regulate IFNβ expression. (A) Immunoblot analysis from THP-1 IFIT-1 cells pre-treated for 16 h with palbociclib and stimulated for the indicated time by cGAMP transfection (500 ng/ml), probed for total and phospho (Ser366) STING, total and phospho (Ser386) IRF3, total and phospho (Tyr701) STAT1 and tubulin. (B) Luciferase activity from THP-1 IFIT-1 cells mock- or pre-treated for 48 h with palbociclib (2 μM) -/+ ruxolitinib (2 μM) and stimulated overnight with IFNβ (1 ng/ml) or by transfection with HT-DNA (50 ng/ml) or cGAMP (500 ng/ml). (C) Luciferase activity from THP-1 Dual Ctrl or IFNAR-/- cells mock- or pre-treated for 48 h with palbociclib (2 μM) and stimulated overnight by transfection with HT-DNA (50 ng/ml) or cGAMP (500 ng/ml). (D) Luciferase activity from THP-1 IFIT-1 cells mock- or pre-treated for 48 h with palbociclib (2 μM) and stimulated overnight with IFNβ (1 ng/ml) or by transfection with cGAMP (500 ng/ml). (E) *IFNβ* expression from THP-1 IFIT-1 cells mock- or pre-treated for 48 h with palbociclib (2 μM) and stimulated for the indicated times by cGAMP transfection (500 ng/ml). (F) *IFNβ* expression from HFFF transduced for 48 h with GFP- or p16INK4a-expressing lentiviruses and stimulated overnight by cGAMP (1000 ng/ml) or HT-DNA (50 ng/ml) transfection. Data are mean ± SD, n = 3, representative of at least 3 repeats. Fold inductions were calculated by normalising each condition with the non-drug treated, non-stimulated (mock) control (B-E) or the FLAG-GFP transduced, non-stimulated (mock) control (F). Statistical analyses were performed using Student’s t-test, with Welch’s correction where appropriate. *P < 0.05, **P < 0.01, ***P < 0.001.

### CDK4/6 negatively regulate IFNβ expression

To gain a mechanistic understanding of how CDK4/6 regulate DNA sensing responses we performed pathway mapping experiments, blotting for the phosphorylated form of key innate signalling molecules downstream of cGAS/STING. Whilst treating THP-1 cells with palbociclib enhanced IFIT-1 reporter activity following cGAMP stimulation as expected (Suppl. Fig 4A), this was not accompanied by enhanced phosphorylation of STING (Ser366) or IRF3 (Ser386) (Fig 3A). This indicates that contrary to previous reported mechanisms(Li *et al.*, 2021; Zhong *et al.*, 2020), CDK4/6 do not inactivate cGAS, and instead regulate innate immunity downstream of these hallmark cytoplasmic signalling events. Consistent with downstream regulation, levels of phosphorylated STAT1 (Tyr701) were increased in palbociclib-treated cells, indicative of CDK4/6-mediated regulation of IFN (Fig 3A). Similar results were observed in primary HFFFs (Suppl. Fig 4B). Total protein expression levels for STING, IRF3 and STAT1 were unaltered following CDK4/6 inhibition (Fig 3A, Suppl. Fig 4B), as was *IRF3* mRNA expression (Suppl. Fig 4C). Palbociclib-dependent enhancement of IFIT-1 reporter activity, which is regulated in an IRF3- and IFN-dependent manner (Mankan *et al.*, 2014), was completely dependent on IFN, as this was abolished in cells treated with JAK inhibitor ruxolitinib (Fig 3B). Furthermore IRF-Luc reporter activity was also not enhanced by palbociclib in THP-1 Dual reporter cells lacking the IFNα/β receptor (IFNAR) (Fig 3C). Increased pSTAT1 could result from increased type I IFN production or increased pathway activation downstream of IFN binding the IFNAR. Palbociclib did not increase ISG expression following direct stimulation of THP-1 cells with IFNβ (Fig 3B, D), ruling out a role for CDK4/6 in regulating the JAK-STAT pathway. Conversely CDK4/6 inhibition did enhance DNA sensing-induced IFNβ mRNA expression in both THP-1 cells (Fig 3E) and HFFF (Fig 3F). Together these findings demonstrate a novel role for CDK4/6 in negatively regulating DNA sensing-induced ISG induction and innate immune activation through limiting type I IFN expression.

### CDK4/6-dependent regulation of DNA sensing is independent of ERVs

Long-term CDK4/6 inhibition in patients and mice was previously reported to correlate with the expression of endogenous retroviral (ERV) elements, consistent with downregulation of the E2F target DNA methyltransferase 1 (DNMT1). ERV upregulation led to an increase in cytoplasmic dsRNA levels, subsequent expression of type III IFN and ISGs, and ultimately enhanced anti-tumour immunity (Goel *et al.*, 2017). CDK4/6-dependent regulation of IFNβ expression however was independent of ERVs as, firstly, neither chemical nor genetic CDK4/6 inhibition in our experiments led to IFNβ or ISG expression in the absence of exogenous stimulation (Fig 1, 2, 3, Suppl. Fig 3) and secondly palbociclib-dependent enhancement of IFIT-1 expression was observed following HT-DNA and cGAMP stimulation in the absence of MAVS (Suppl. Fig 5A), the major RNA sensing adaptor protein that is important for the detection of ERV RNA(Chiappinelli *et al*, 2015; Roulois *et al*, 2015; Tie *et al*, 2018). As a control, MAVS-/- cells did not respond to RNA sensing agonist poly I:C (Suppl. Fig 5A). Furthermore, cGAMP-induced reporter activity was also increased in palbociclib-treated cells lacking cGAS (Suppl. Fig 5A), which has also been implicated in the sensing of ERVs (Lima-Junior *et al*, 2021). As expected, cGAS-/- cells failed to respond to HT-DNA (Suppl. Fig 5A). These data imply a direct role for CDK4/6 in negatively regulating type I IFN expression, independent of E2F/DNMT1-mediated upregulation of ERV elements.

### CDK4/6 regulate type I IFN downstream of multiple PRRs

Given that CDK4/6 regulation of innate immunity mapped to the level of IFNβ expression, we questioned whether pathways other than DNA sensing may be regulated in this manner. To this end we treated THP-1 cells with palbociclib and stimulated them with other agonists of type I IFN expression including RNA sensing agonists transfected poly I:C and Sendai virus or TLR4 agonist lipopolysaccharide (LPS) and found that IFIT-1 reporter activity (Fig 4A), endogenous ISG expression (Fig 4B) and CXCL-10 protein secretion (Fig 4C) were all significantly enhanced by CDK4/6 inhibition. The same was observed in HFFF transfected with poly I:C (Fig 4D) and following p16INK4a overexpression (Fig 4E). Co-treatment with ruxolitinib demonstrated that augmented ISG expression was again entirely dependent on IFN (Suppl. Fig A) and enhanced IFNβ transcription was observed in conditions of CDK4/6 inhibition (Fig 4F). Increased innate responses following CDK4/6 inhibition was again independent of ERV sensing as this was independent of cGAS (Suppl. Fig 6B) and MAVS (Suppl. Fig. 6C) expression. Taken together these data reveal a broad role for cell cycle regulatory proteins CDK4/6 in negatively regulating type I IFN and subsequent ISG expression downstream of multiple PRRs, including DNA sensing, RNA sensing and TLR pathways.

**Fig 4:**
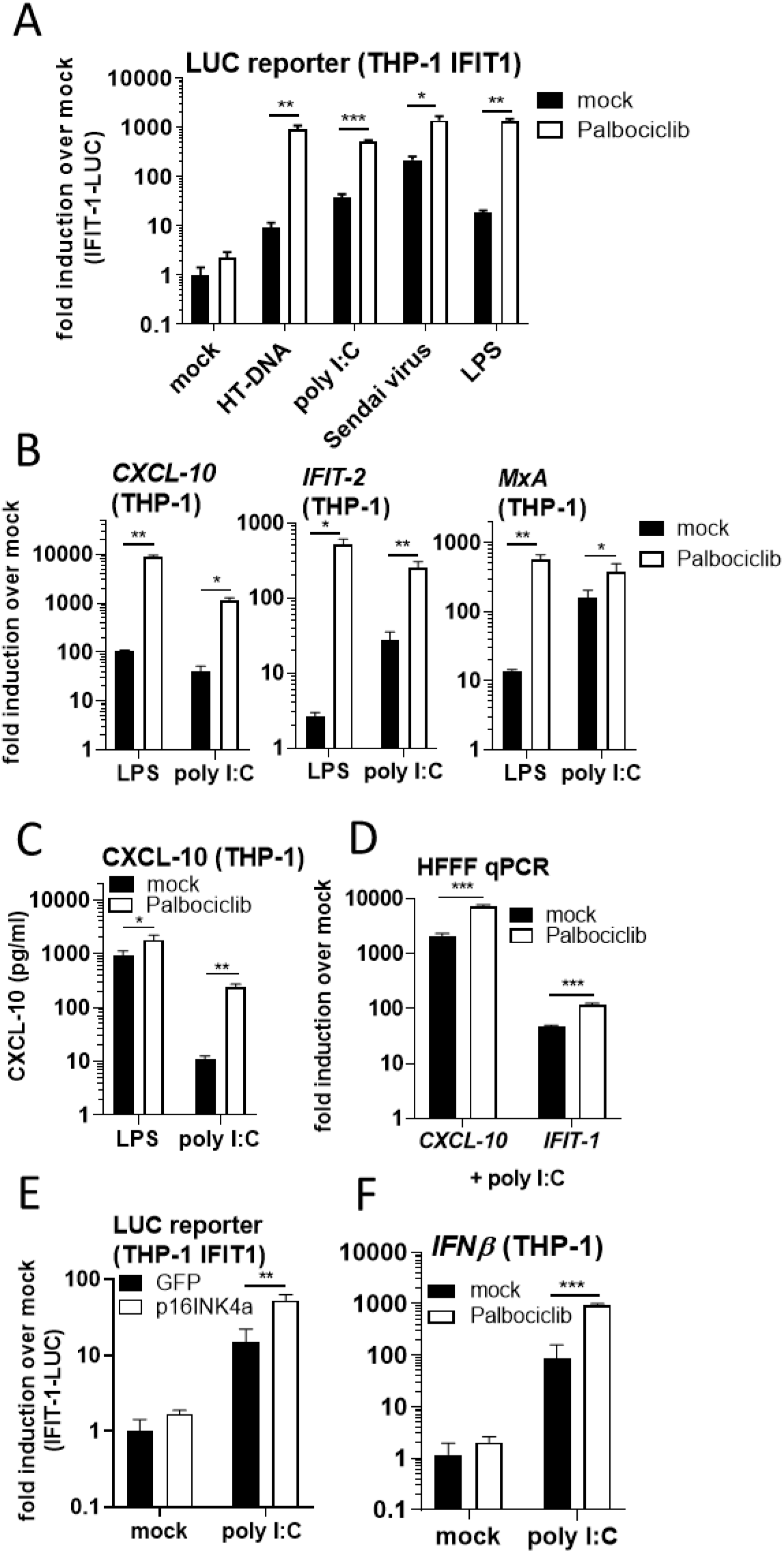
CDK4/6 regulate type I IFN downstream of multiple PRRs. Luciferase activity from THP-1 IFIT-1 cells mock- or pre-treated for 48 h with palbociclib (2 μM) and stimulated overnight with LPS (100 ng/ml), Sendai virus infection (0.2 HA U/ml) or by HT-DNA (20 ng/ml), cGAMP (500 ng/ml) or poly I:C (500 ng/ml) transfection. (B) ISG qRT-PCR from THP-1 IFIT-1 cells mock- or pre-treated for 48 h with palbociclib (2 μM) and stimulated overnight with LPS (100 ng/ml) or by poly I:C (500 ng/ml) transfection. (C) ELISA from (B). (D) ISG qRT-PCR from HFFF mock- or pre-treated for 48 h with palbociclib (2 μM) and stimulated overnight by poly I:C (200 ng/ml) transfection. (E) Luciferase activity of THP-1 IFIT-1 cells transduced for 48 h with FLAG-GFP- or FLAG-p16INK4a-expressing lentiviruses and stimulated overnight by poly I:C (500 ng/ml) transfection. (F) *IFNβ* expression from THP-1 IFIT-1 cells mock- or pre-treated for 48 h with palbociclib (2 μM) and stimulated for 4 h by poly I:C transfection (500 ng/ml). Data are mean ± SD, n = 3, representative of at least 3 repeats. Fold inductions were calculated by normalising each condition with the non-drug treated, non-stimulated (mock) control (A-D, F), or with FLAG-GFP lentivirus transduced, non-stimulated (mock) control (E). Statistical analyses were performed using Student’s t-test, with Welch’s correction where appropriate. *P < 0.05, **P < 0.01, ***P < 0.001.

## Discussion

Innate immune sensing must be tightly regulated to avoid aberrant recognition of cellular patterns and the induction of autoimmune conditions. Here we present a novel mechanism of such regulation dependent on cell cycle regulatory proteins CDK4/6. Manipulation of CDK4/6 by chemical (Fig 1) or genetic inhibition (Fig 2), or by shRNA-mediated depletion (Fig 2) significantly increased DNA sensing responses in both primary and transformed cells of human and murine origin (Suppl. Fig 3). This regulation mapped downstream of IRF3 phosphorylation at the level of IFNβ expression, resulting in increased STAT1 phosphorylation and enhanced IFN-dependent ISG expression (Fig 3). Consistent with these findings, CDK4/6 also negatively regulated innate immune activation downstream of other PRR pathways that induce type I IFN expression, including RNA and TLR sensing (Fig 4). This regulation was independent of cell cycle arrest (Suppl. Fig 1), senescence (Suppl. Fig 1) and ERV upregulation (Suppl. Fig 5), all of which have been observed following long-term CDK4/6 inhibitor treatment and result in inflammatory cytokine expression(Gluck *et al.*, 2017; Goel *et al.*, 2017; Yoshida *et al.*, 2016). Together these data reveal a novel and direct role for CDK4/6 in dampening type I IFN responses.

In this study we mapped CDK4/6-dependent regulation of ISG expression to the level of IFNβ expression. As cytoplasmic signalling events such as phosphorylation of STING, and phosphorylation of IRF3 at residue Ser386, which is necessary for nuclear translocation(Chen *et al*, 2008), were not enhanced following CDK4/6 inhibition, this implies a potential role for these kinases in regulating IFN expression at the promoter level. This could, for example, be mediated by enhanced transcription factor or positive regulator binding, reduced binding of repressors and/or epigenetic changes that result in a shift from a repressive to an active chromatin state. Indeed, methyltransferase DNMT1 is an E2F target gene(Kimura *et al*, 2003) and whether its expression following CDK4/6 inhibition contributes to enhanced IFN expression will be important future work. Also, whether canonical substrate pRb and the downstream E2Fs, which are necessary for CDK4/6-dependent G1 to S phase transition, play a role in innate immune regulation, or whether it involves a non-canonical substrate, of which a number have already been described(Jirawatnotai *et al*, 2014), remains to be determined. CDK4/6 inhibitors target the ATP-binding domains of CDK4 and 6 (Toogood *et al*, 2005), suggesting their kinase activity is likely required for IFN regulation. Interestingly the expression of IRF3 itself can be directly repressed by E2F1(Xu *et al*, 2011), however we did not observe enhanced levels of IRF3 following palbociclib treatment up to 48 h, either at the transcript (Suppl. Fig 4C) or protein level (Fig 3A, Suppl. Fig. 4B). Uncovering the CDK4/6 target(s) that is responsible for the described phenotype may reveal novel mechanisms regulating IFNβ expression, which could be manipulated for therapeutic benefit.

Palbociclib and other CDK4/6 inhibitors are approved for the treatment of breast cancer and have been shown in patients to boost tumour immunity(Goel *et al.*, 2017). What is unclear however is whether this increased immunogenicity is a direct consequence of innate immune activation. CDK4/6 inhibition augments innate responses by multiple mechanisms, including increased type III IFN expression through reactivation of ERV elements (Goel *et al.*, 2017), increased type II IFN, partially dependent on STING(Liu *et al*, 2021), induction of senescence and the ensuing senescence-associated secretory phenotype(Gluck *et al.*, 2017; Yoshida *et al.*, 2016) and here, enhanced PAMP-induced type I IFN expression. Together these may increase CDK4/6 inhibitor treatment efficacy *in vivo* through increased immune cell recruitment, antigen presentation and adaptive immune responses.

Interestingly, roles for other CDKs in regulating innate immunity have previously been described. Cingoz et al observed that some CDKs were necessary for type I IFN induction, although this was not true for CDK4/6 (Cingoz & Goff, 2018). Furthermore there have been reports of a pan-CDK inhibitor suppressing TLR signalling (Zoubir *et al*, 2011) and inhibition of CDK2 activity resulting in enhanced NF-κB-dependent gene expression (Perkins *et al*, 1997). These findings reveal a complex interplay between cell division and the host innate immune response.

Aberrant cGAS/STING activation has been linked to multiple autoimmune disorders including Aicardi-Goutieres syndrome(Gray *et al*, 2015), systemic lupus erythematosus(Kato *et al*, 2018) and rheumatoid arthritis(Wang *et al*, 2019), therefore tight regulation of this pathway during homeostasis is critical to avoid disease. How cells tolerate nuclear membrane dissolution during mitosis, for example, without activating a detrimental IFN response remains an important question. Recent studies have demonstrated that cGAS itself is regulated in multiple ways during cellular replication. For example, cGAS associates with chromatin following nuclear envelope breakdown, preventing its oligomerisation and subsequent activation(Li *et al.*, 2021). Furthermore cGAS is inactivated through phosphorylation by mitotic kinases such as Aurora kinase B(Li *et al.*, 2021) and CDK1(Zhong *et al.*, 2020), both of which block cGAMP production. Barrier-to-autointegration factor 1, which binds chromatin and is essential for nuclear membrane reformation, has also been shown to compete with cGAS for DNA binding following nuclear envelope dissolution, again restricting cGAS activity during mitosis(Guey *et al*, 2020). Here we have discovered a broader mechanism dampening innate activation during cell division dependent on CDK4/6 restriction of IFNβ expression, occurring downstream of cGAS/STING and other PRRs. Our findings reveal innate immune regulation during earlier stages of cell division than have previously been reported, which may be important for establishing an immune suppressed state in preparation for DNA replication during S phase. How long CDK4/6-induced suppression of type I IFN continues when these kinases are no longer active remains to be tested, but it is conceivable that any changes CDK4/6 induce at the IFNβ promoter may continue to dampen expression through subsequent cell cycle stages. Together, these findings define cell division as an innate immune privileged process, expanding our understanding of how cells regulate IFN production during homeostasis and avoid autoimmune reaction.

## Acknowledgments

We thank Veit Hornung for kindly providing THP-1-IFIT-1 cells. RPS and CM were funded by a collaborative MRC award (MRC_PC_19052) and the University of Surrey Faculty Research Support Funds. CM was also funded by the BBSRC (BB/T006501/1, BB/V015265/1). GJT was funded through a Wellcome Trust Senior Biomedical Research Fellowship (108183) followed by a Wellcome Investigator Award (220863), the European Research Council under the European Union’s Seventh Framework Programme (FP7/2007-2013)/ERC (grant HIVInnate 339223) a Wellcome Trust Collaborative award (214344) and the National Institute for Health Research University College London Hospitals Biomedical Research Centre.

## Author Contributions

RPS conceived the study. RPS, AE, SL and HA performed the experiments and analysed the data. RPS, GJT and CM wrote the manuscript and obtained funding.

## Methods

### Cells and reagents

HEK293T, human foetal foreskin fibroblasts (HFFF), A549 and U2OS cells were maintained in DMEM (Gibco) supplemented with 10 % foetal bovine serum (FBS, Labtech) and 100 U/ml penicillin plus 100 μg/ml streptomycin (Pen/Strep; Gibco). Murine McCoy cells were maintained in MEM (Gibco) supplemented with 10 % FBS and Pen/Strep. THP-1 cells were maintained in RPMI (Gibco) supplemented with 10 % FBS and Pen/Strep. THP-1-IFIT-1 cells that had been modified to express Gaussia luciferase under the control of the *IFIT-1* promoter were described previously (Mankan *et al.*, 2014). THP-1-IFIT-1 MAVS-/- cells were also previously described(Sumner *et al*, 2020; Tie *et al.*, 2018). THP-1 Dual Control, IFNAR-/- and cGAS-/- cells were obtained from Invivogen. Inhibitors against CDK1 (RO-3306), CDK2 (K03861) and CDK4/6 (palbociclib) and piperine were all from Sigma. CDK4/6 inhibitor ribociclib was obtained from Cambridge Biosciences. Lovastatin was from Abcam. JAK inhibitor ruxolitinib was from CELL guidance systems. Lipopolysaccharide and IFNβ were obtained from Peprotech. Sendai virus was from Charles River Laboratories. Herring-testis DNA was obtained from Sigma. cGAMP and poly I:C were from Invivogen. For stimulation of cells by transfection, transfection mixes were prepared using lipofectamine 2000 according to the manufacturer’s instructions (Invitrogen).

### Plasmids

The p16INK4a sequence was amplified from THP-1 cDNA using forward (5’-GAATGCGGCCGCGGAGCCGGCGGCGGGGAGC-3’) and reverse (5’-GCCTCTAGACTCGAGTCAATCGGGGATGTCTGAG-3’) primers and cloned into a pcDNA4/TO FLAG (N-terminal tag) vector using NotI and XbaI. FLAG-tagged p16INK4a was then excised from this vector with BamHI and XhoI and cloned into lentiviral genome plasmid SFXUP (expressing p16INK4a under the control of the spleen focus-forming virus (SFFV) promoter and puromycin under the control of the ubiquitin promoter). SFXUP expressing FLAG-GFP served as a control and was used previously (Georgana & Maluquer de Motes, 2019).

For transient depletion of CDKs by shRNA, annealed oligos were cloned with BamHI and EcoRI into HIV-1-based shRNA expression vector HIVSiren (Schaller *et al*, 2011). The shCtrl sequence has been previously described(Fletcher *et al*, 2015).

CDK1 shRNA top: 5’-GATCCGGGATTCCAGGTTATATCTCTTCAAGAGAGAGATATAACCTGGAATCCTTTTTTG-3’

CDK1 shRNA bottom: 5’-AATTCAAAAAAGGATTCCAGGTTATATCTCTCTCTTGAAGAGATATAACCTGGAATCCCG-3’

CDK2 shRNA top: 5’-GATCCGAGGTTATATCCAATAGTAGTTCAAGAGACTACTATTGGATATAACCTTTTTTTG-3’

CDK2 shRNA bottom: 5’-AATTCAAAAAAAGGTTATATCCAATAGTAGTCTCTTGAACTACTATTGGATATAACCTCG-3’

CDK4 shRNA top: 5’-GATCCGAGGCCTAGATTTCCTTCATTTCAAGAGAATGAAGGAAATCTAGGCCTTTTTTTG-3’

CDK4 shRNA bottom: 5’-AATTCAAAAAAAGGCCTAGATTTCCTTCATTCTCTTGAAATGAAGGAAATCTAGGCCTCG-3’

CDK6 shRNA top: 5’-GATCCGGTTCAGATGTTGATCAACTTTCAAGAGAAGTTGATCAACATCTGAACTTTTTTG-3’

CDK6 shRNA bottom: 5’-AATTCAAAAAAGTTCAGATGTTGATCAACTTCTCTTGAAAGTTGATCAACATCTGAACCG-3’

### PI staining

One million cells were pelleted and washed once with PBS. Cells were fixed in cold 70 % ethanol and stored at 4 °C until staining. To stain for DNA content, cells were washed twice in PBS and the pellet then treated with RNase A diluted in PBS (100 μg/ml, Sigma) before incubation with propidium iodide (50 μg/ml, diluted in PBS, Sigma). Cells were analysed by flow cytometry using the FACS Calibur (BD) and data analysis was performed using FlowJo software.

### Production of lentiviral particles in 293T cells

Lentiviral particles were produced by transfecting 10 cm dishes of HEK293T cells with 1.5 μg of genome plasmid (HIVSiren for shRNA, or SFXUP for GFP- or p16INK4a-expressing vectors), 1 μg of p8.91 packaging plasmid (Zufferey *et al*, 1997), and 1 μg of vesicular stomatitis virus-G glycoprotein expressing plasmid pMDG (Genscript) using Fugene 6 transfection reagent (Promega) according to the manufacturer’s instructions. Virus supernatants were harvested at 48 and 72 h post-transfection, pooled and stored at -80 °C. THP-1 cells were transduced by spinoculation (1000 *xg*, 1 h, room temperature) and adherent HFFFs were transduced by incubation with lentivector-containing medium. Cells were incubated for 48 h without drug selection to allow depletion or overexpression of target genes before being used for experiments.

### Luciferase reporter assays

Following pre-treatment of cells with inhibitors, or lentiviral transduction for depletion/overexpression, THP-1 cells were counted, plated in 96 well plates and stimulated with the indicated agonists at 2×10^5^ cells/ml. Gaussia/Lucia luciferase activity was measured 16-24 h later by transferring 10 μl supernatant to a white 96 well assay plate, injecting 50 μl per well of coelenterazine substrate (Nanolight Technologies, 2 μg/ml) and analysing luminescence on a CLARIOstar luminometer (BMG). Data were normalised to a mock, non-drug-treated control to generate a fold induction.

### ISG qPCR

RNA was extracted from cells using a total RNA purification kit (QIAGEN) according to the manufacturer’s protocol. Five hundred ng RNA was used to synthesise cDNA using Superscript III reverse transcriptase (Invitrogen), also according to the manufacturer’s protocol. For IFNβ qPCR, RNA was treated with DNase prior to cDNA synthesis (Invitrogen). cDNA was diluted 1:5 in water and 2 μl used as a template for real-time PCR using SYBR® Green PCR master mix (Applied Biosystems) and a Quant Studio 5 (Applied Biosystems) or LightCycler 96 (Roche) real-time PCR machine. Expression of each gene was normalised to an internal control (*GAPDH* or *HPRT*) and these values were then normalised to mock, non-drug treated control cells to yield a fold induction. The following primers were used:

Human:

*GAPDH* Fwd 5’-GGGAAACTGTGGCGTGAT-3’

*GAPDH* Rev 5’-GGAGGAGTGGGTGTCGCTGTT-3’

*CXCL-10* Fwd 5’-TGGCATTCAAGGAGTACCTC-3’,

*CXCL-10* Rev 5’-TTGTAGCAATGATCTCAACACG-3’

*IFIT-1* Fwd 5’-CCTCCTTGGGTTCGTCTACA-3’

*IFIT-1* Rev 5’-GGCTGATATCTGGGTGCCTA-3’

*IFIT-2* Fwd 5’-CAGCTGAGAATTGCACTGCAA-3’

*IFIT-2* Rev 5’-CGTAGGCTGCTCTCCAAGGA-3’

*MxA* Fwd 5’-ATCCTGGGATTTTGGGGCTT-3’

*MxA* Rev 5’-CCGCTTGTCGCTGGTGTCG-3’

*2’5’OAS* Fwd 5’-TGTGTGTGTCCAAGGTGGTA-3’

*2’5’OAS* Rev 5’-TGATCCTGAAAAGTGGTGAGAG-3’

*IFNβ* Fwd 5’-ACATCCCTGAGGAGATTAAGCA-3’

*IFNβ* Rev 5’-GCCAGGAGGTTCTCAACAATAG-3’

Mouse:

*HPRT* Fwd 5’-GGTTAAGCAGTACAGCCCCAA-3’

*HPRT* Rev 5’-ATAGGCACATAGTGCAAATCA-3’

*CXCL-10* Fwd 5’-ACTGCATCCATATCGATGAC-3’

*CXCL-10* Rev 5’-TTCATCGTGGCAATGATCTC-3’

*IFIT-1* Fwd 5’-ACCATGGGAGAGAATGCTGAT-3’

*IFIT-1* Rev 5’-GCCAGGAGGTTGTGC-3’

*2’5’OAS* Fwd 5’-TGAGCGCCCCCCATCT-3’

*2’5’OAS* Rev 5’-CATGACCCAGGACATCAAAGG-3’

### ELISA

Cell supernatants were harvested for ELISA at 24 h post-infection/stimulation and stored at - 80 °C. CXCL-10 protein was measured using Duoset ELISA reagents (R&D Biosystems) according to the manufacturer’s instructions.

### Immunoblotting

For immunoblot analysis, THP-1 (3×10^6^ cells) or HFFF (in 12 well plates) cells were lysed in a cell lysis buffer containing 50 mM Tris pH 8, 150 mM NaCl, 0.1 % (w/v) sodium dodecyl sulfate (SDS), 0.5 % (w/v) sodium deoxycholate, 1 % (v/v) NP-40 supplemented with protease inhibitors (Roche), clarified by centrifugation at 14,000 x *g* for 10 min and boiled in 6X Laemmli buffer for 5 min. For phosphoblotting experiments phosphatase inhibitors (Roche) were included in the lysis buffer and samples were analysed immediately. Proteins were separated by SDS-PAGE on 12 % polyacrylamide gels. After PAGE, proteins were transferred to a Hybond ECL membrane (Amersham biosciences) using a semi-dry transfer system (Biorad). Primary antibodies were from the following sources: mouse anti-β-actin (Abcam), mouse-anti-MCM2 (BD), mouse-anti-FLAG (Sigma), mouse-anti-tubulin (EMD Millipore), rabbit-anti-CDK1 (Cell Signaling), rabbit-anti-CDK2 (Cell Signaling), rabbit-anti-CDK4 (Cell Signaling), mouse-anti-CDK6 (Cell Signaling), rabbit-anti-STING (Cell Signaling), rabbit-anti-pSTING (Ser 366, Cell Signaling), rabbit-anti-IRF3 (Abcam), rabbit-anti-pIRF3 (Ser 386, Abcam), rabbit-anti-STAT1 (Cell Signaling), rabbit-anti-pSTAT1 (Tyr 701, Cell Signaling). Primary antibodies were detected with goat-anti-mouse/rabbit IRdye 680/800 infrared dye secondary antibodies and membranes imaged using an Odyssey Infrared Imager (LI-COR Biosciences).

### Statistical analyses

Statistical analyses were performed using an unpaired Student’s t-test (with Welch’s correction where variances were unequal). * *P*<0.05, ** *P*<0.01, *** *P*<0.001.

## Figures

**Suppl Fig 1:**
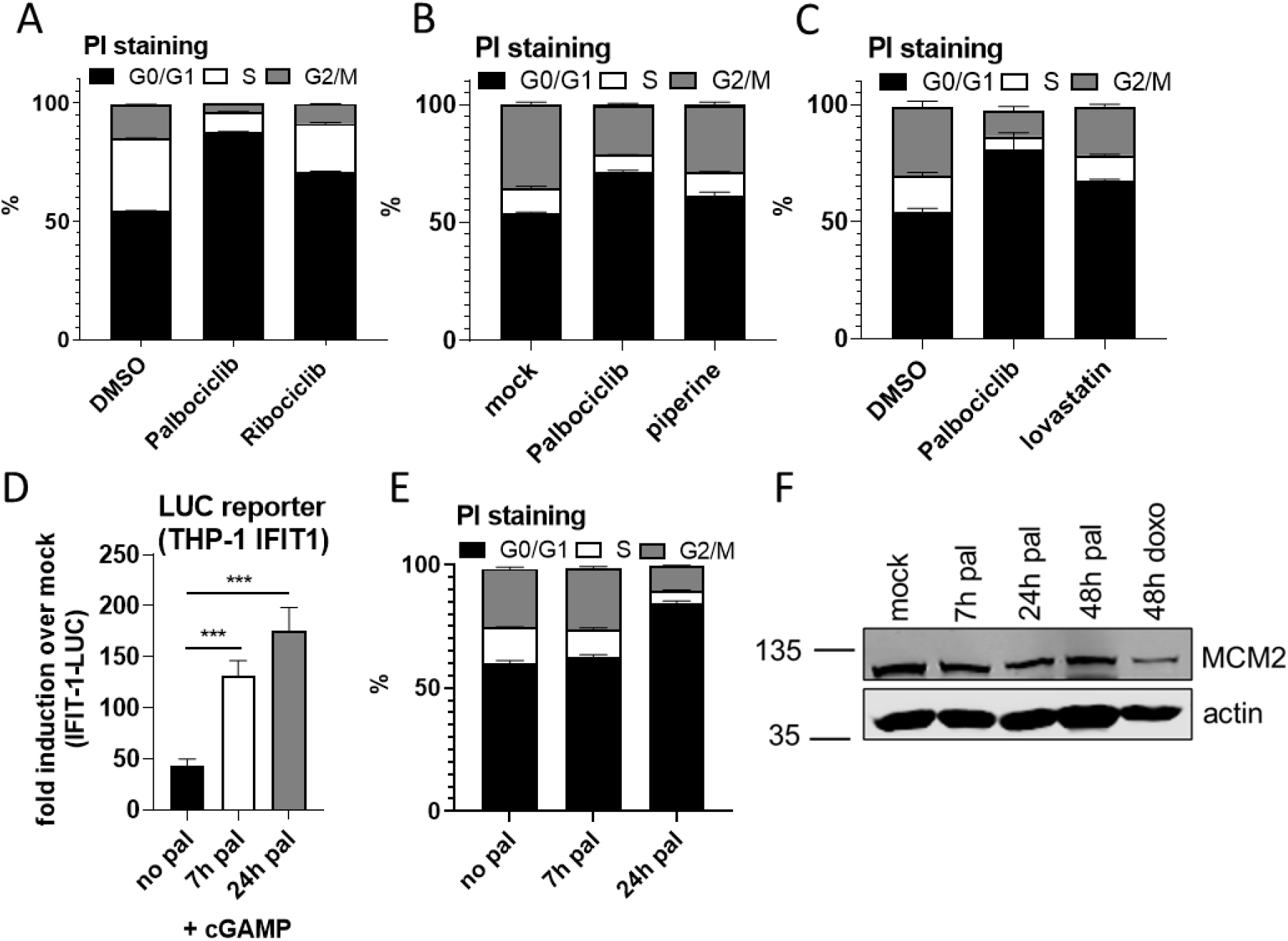
CDK4/6 inhibitor-mediated regulation of DNA sensing is independent of cell cycle arrest and senescence. (A) PI staining from Fig 1E. (B) PI staining from Fig 1F. (C) PI staining from Fig 1G. (D) Luciferase activity from THP-1 IFIT-1 cells mock-treated, or pre-treated with palbociclib (2 μM) for 7 or 24 h and then stimulated overnight by transfection with cGAMP (500 ng/ml). (E) PI staining from (D). (F) Immunoblot of THP-1 IFIT-1 cells treated for the indicated times with palbociclib (2 μM) or doxorubicin (1 μM) and probed for MCM2 and β-actin. Data are mean ± SD, n = 3, representative of at least 3 repeats. Fold inductions were calculated by normalising each condition with the non-drug treated, non-stimulated (mock) control. Statistical analyses were performed using Student’s t-test, with Welch’s correction where appropriate. ***P < 0.001.

**Suppl. Fig 2:**
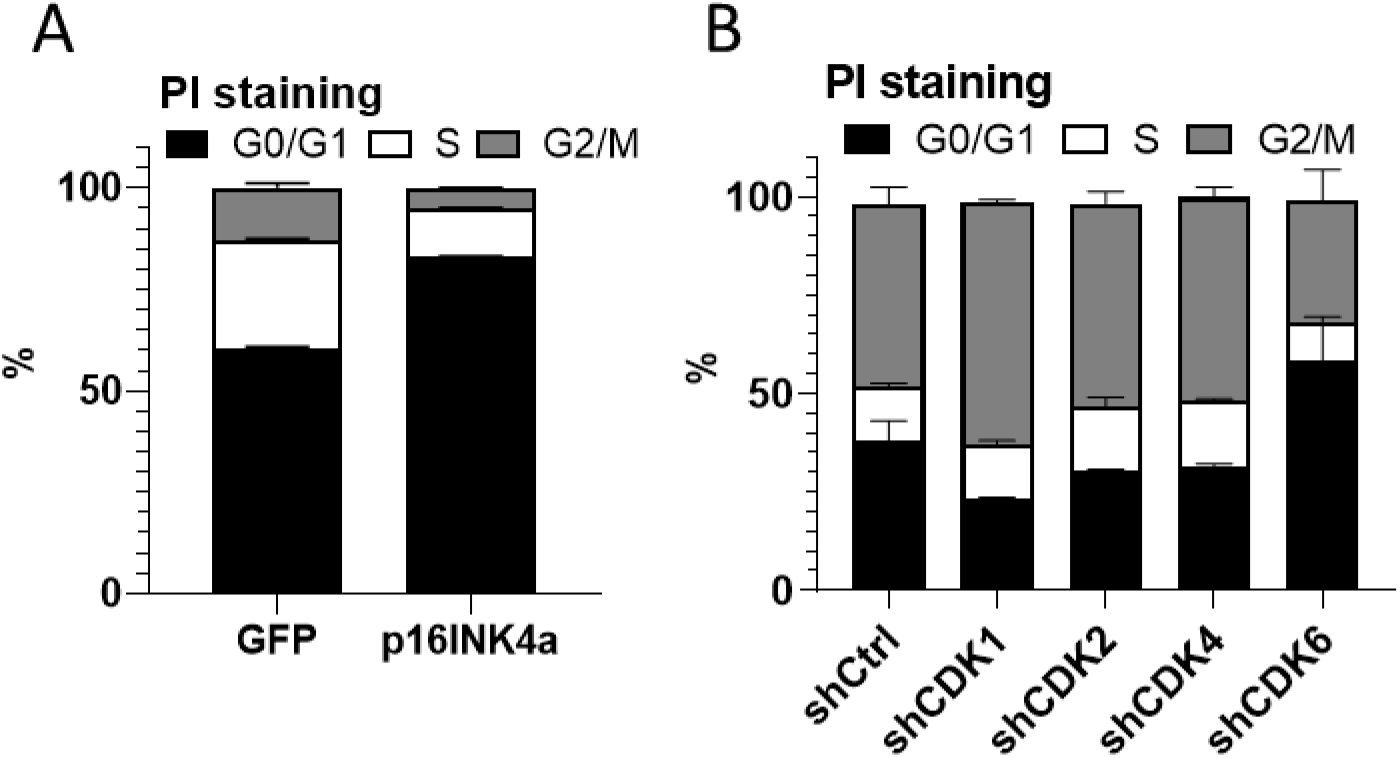
CDK4/6 negatively regulate DNA sensing. (A) PI staining from Fig 2A, B. (B) PI staining from Fig 2C, D. Data are mean ± SD, n = 3, representative of at least 3 repeats.

**Suppl. Fig 3:**
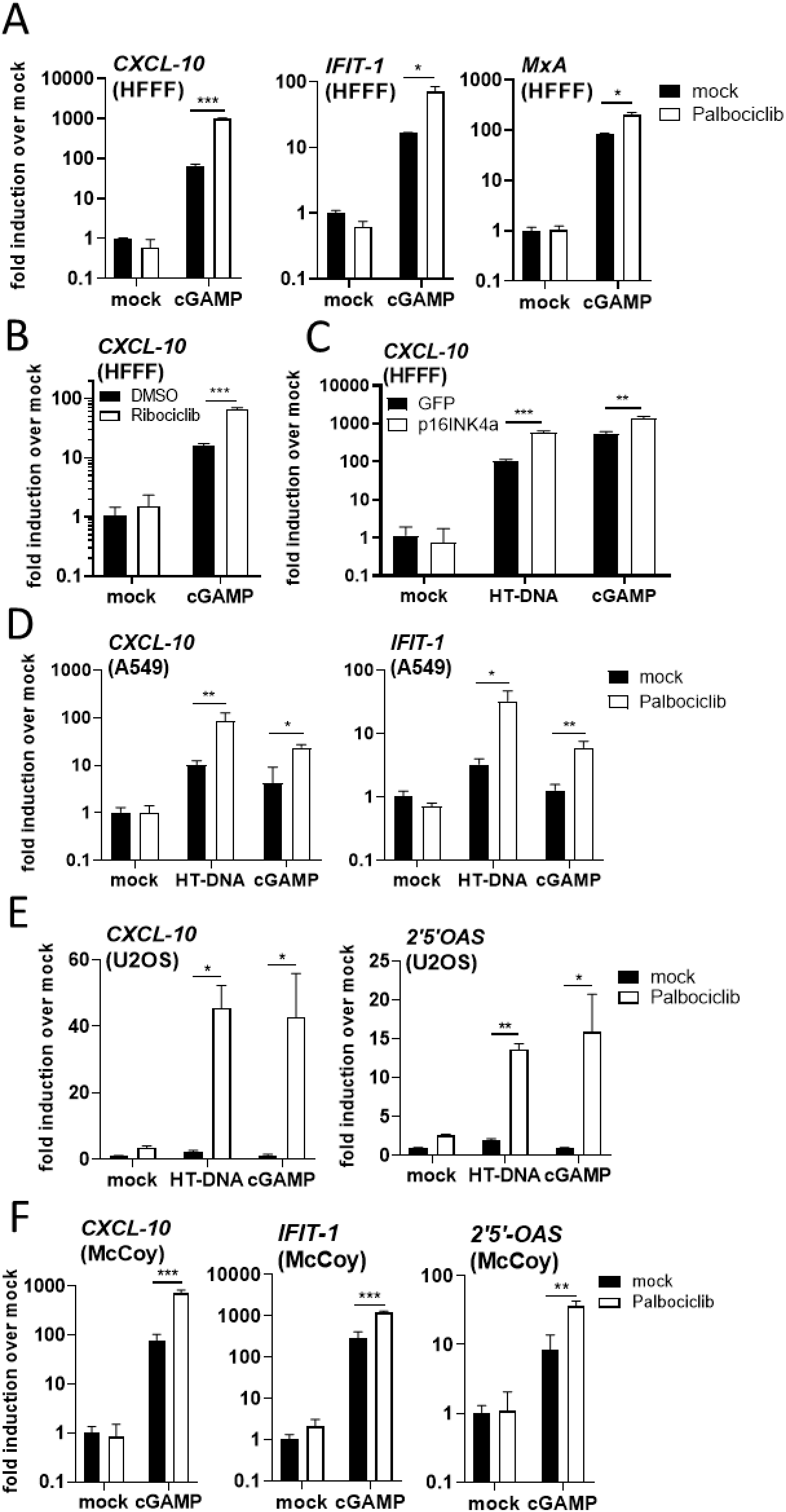
CDK4/6 negatively regulate DNA sensing. (A, B) ISG qRT-PCR from HFFFs cells pre-treated for 48 h with palbociclib (2 μM, A) or ribociclib (2 μM, B) and stimulated overnight by transfection with cGAMP (1000 ng/ml). (C) ISG qRT-PCR from HFFFs transduced with FLAG-GFP/INK4a-expressing lentivirus for 48 h and then stimulated overnight by transfection with HT-DNA (50 ng/ml) or cGAMP (1000 ng/ml). (D, E) ISG qRT-PCR from A549 (D) or U2OS (E) cells pre-treated for 48 h with palbociclib (2 μM) and stimulated by overnight transfection with HT-DNA (100 ng/ml) or cGAMP (1000 ng/ml). (F) ISG qRT-PCR from murine McCoy fibroblasts pre-treated for 48 h with palbociclib (2 μM) and stimulated by overnight transfection with cGAMP (2000 ng/ml). Data are mean ± SD, n = 3, representative of at least 3 repeats. Fold inductions were calculated by normalising each condition with the non-drug treated, non-stimulated (mock) control (A, B, D-F) or the FLAG-GFP transduced, non-stimulated (mock) control (C). Statistical analyses were performed using Student’s t-test, with Welch’s correction where appropriate. *P < 0.05, **P < 0.01, ***P < 0.001.

**Suppl. Fig 4:**
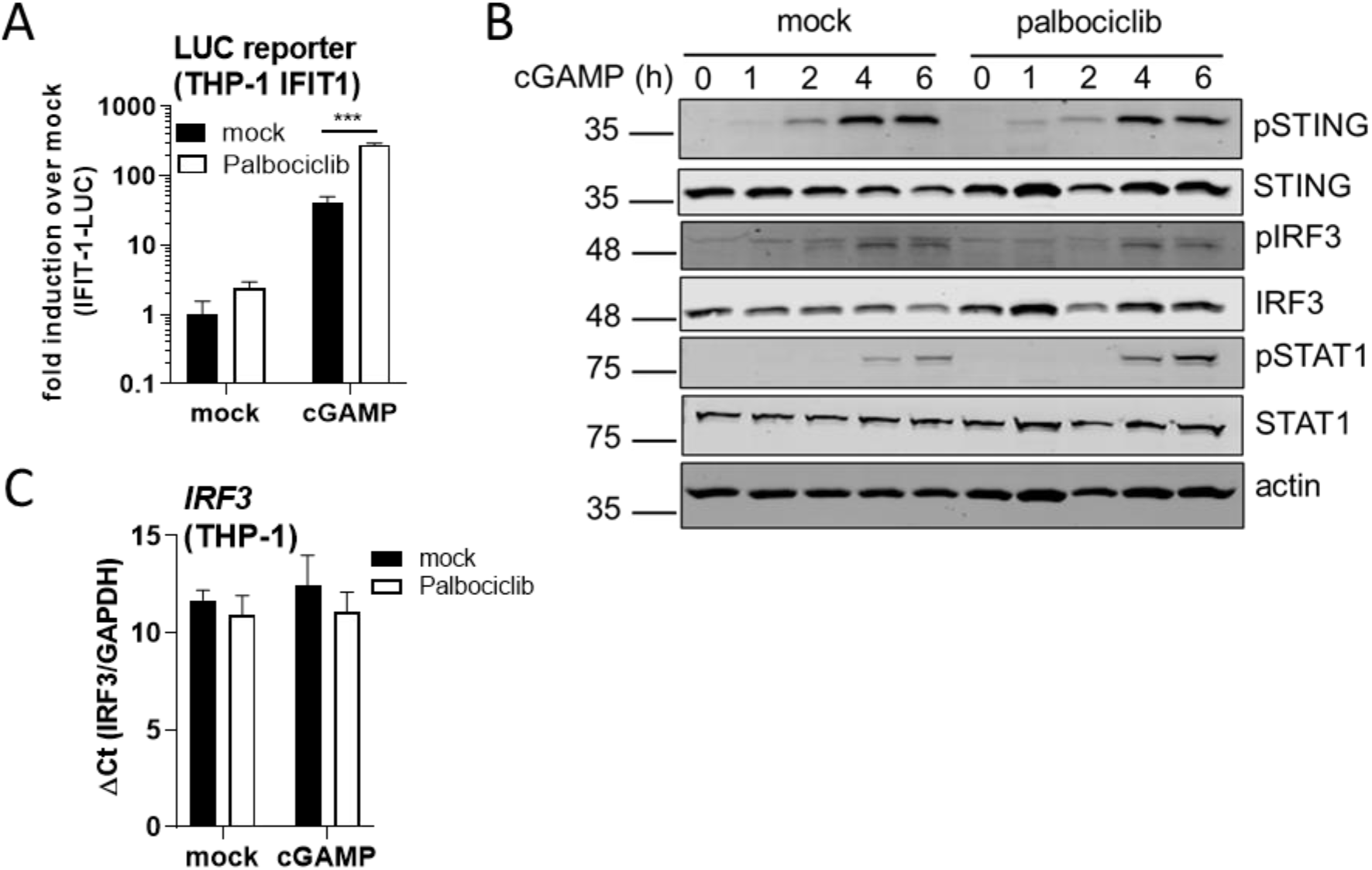
CDK4/6 negatively regulate IFNβ expression. (A) Luciferase activity from Fig 3A. (B) Immunoblot analysis from HFFF cells pre-treated for 16 h with palbociclib and stimulated for the indicated time by cGAMP transfection (1000 ng/ml), probed for total and phospho (Ser366) STING, total and phospho (Ser386) IRF3, total and phospho (Tyr701) STAT1 and tubulin. (C) *IRF3* expression from THP-1 IFIT-1 cells mock- or pre-treated for 48 h with palbociclib (2 μM) and stimulated overnight by cGAMP transfection (500 ng/ml). Data are mean ± SD, n = 3, representative of at least 3 repeats. Fold inductions were calculated by normalising each condition with the non-drug treated, non-stimulated (mock) control (B). ΔCt calculated by normalising to *GAPDH* expression (C). Statistical analyses were performed using Student’s t-test, with Welch’s correction where appropriate. ***P < 0.001.

**Suppl Fig 5:**
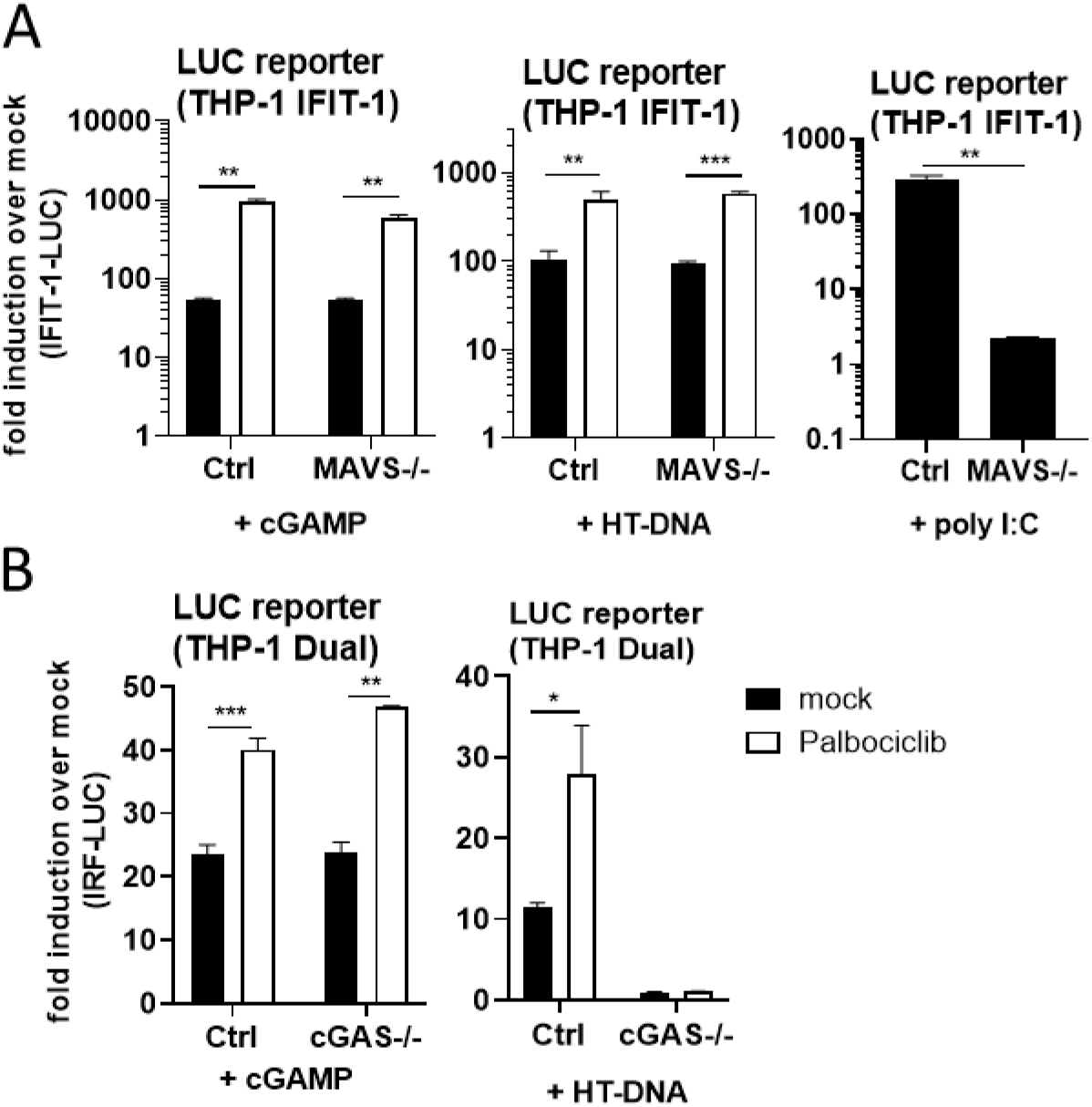
CDK4/6-dependent regulation of ISGs is independent of ERV sensing. (A) Luciferase activity from THP-1 IFIT-1 Ctrl or MAVS-/- cells mock- or pre-treated for 48 h with palbociclib (2 μM) and stimulated overnight by HT-DNA (50 ng/ml), cGAMP (500 ng/ml) or poly I:C (125 ng/ml) transfection. (B) Luciferase activity from THP-1 Dual Ctrl or cGAS-/- cells mock- or pre-treated for 48 h with palbociclib (2 μM) and stimulated overnight by HT-DNA (50 ng/ml) or cGAMP (500 ng/ml) transfection. Data are mean ± SD, n = 3, representative of at least 3 repeats. Fold inductions were calculated by normalising each condition with the non-drug treated, non-stimulated (mock) control. Statistical analyses were performed using Student’s t-test, with Welch’s correction where appropriate. **P < 0.01, ***P < 0.001.

**Suppl Fig 6:**
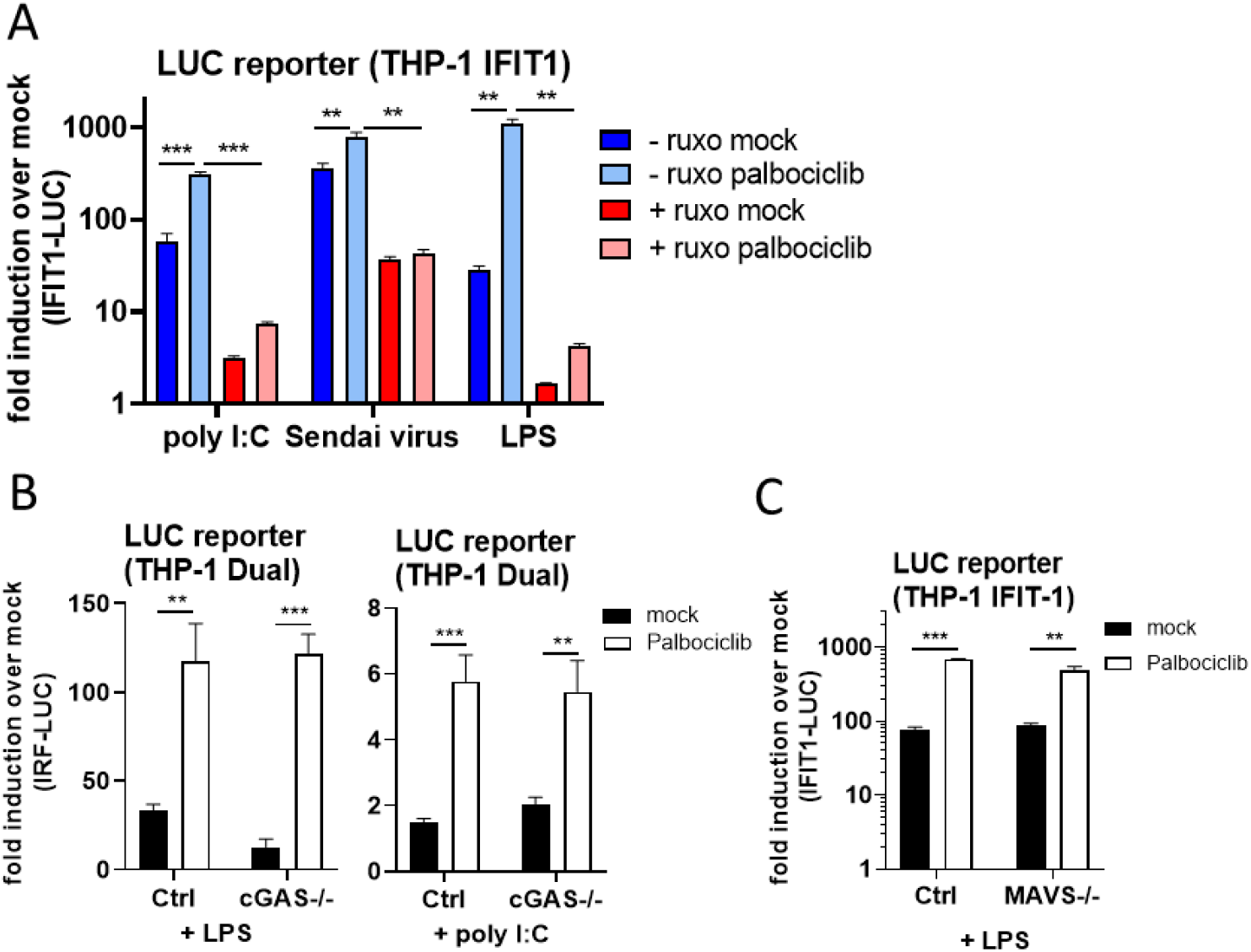
CDK4/6 regulate type I IFN downstream of multiple PRRs. (A) Luciferase activity from THP-1 IFIT-1 cells mock- or pre-treated for 48 h with palbociclib (2 μM) -/+ ruxolitinib (2 μM) and stimulated overnight with LPS (100 ng/ml), infection with Sendai virus (0.2 HA U/ml) or by transfection with poly I:C (500 ng/ml). (B) Luciferase activity from THP-1 Dual Ctrl or cGAS-/- cells mock- or pre-treated for 48 h with palbociclib (2 μM) and stimulated overnight with LPS (100 ng/ml) or by poly I:C (1000 ng/ml) transfection. (C) Luciferase activity from THP-1 IFIT-1 Ctrl or MAVS-/- cells mock- or pre-treated for 48 h with palbociclib (2 μM) and stimulated overnight with LPS (100 ng/ml). Data are mean ± SD, n = 3, representative of at least 3 repeats. Fold inductions were calculated by normalising each condition with the non-drug treated, non-stimulated (mock) control. Statistical analyses were performed using Student’s t-test, with Welch’s correction where appropriate. **P < 0.01, ***P < 0.001.

